# Epitope-Evaluator: an interactive web application to study predicted T-cell epitopes

**DOI:** 10.1101/2022.05.09.491119

**Authors:** Luis Fernando Soto, David Requena, Juan Ignacio Fuxman Bass

## Abstract

Multiple immunoinformatic tools have been developed to predict T-cell epitopes from protein amino acid sequences for different major histocompatibility complex (MHC) alleles. These prediction tools output hundreds of potential peptide candidates which require further processing; however, these tools are either not graphical or not friendly for non-programming users. We present Epitope-Evaluator, a web tool developed in the Shiny/R framework to interactively analyze predicted T-cell epitopes. This includes providing the distribution of epitopes across a selected set of MHC alleles, the promiscuity and conservation of epitopes, and their density and location within antigens. Epitope-Evaluator requires as input the fasta file of protein sequences and the output prediction file coming out from any predictor. By choosing different cutoffs and parameters, users can produce several interactive plots and tables that can be downloaded as JPG and text files, respectively. Epitope-Evaluator removes the programming barrier and provides intuitive tools, allowing a straightforward interpretation and graphical representations that facilitate the selection of candidate epitopes for experimental evaluation.

**Author Summary:** With the advent of the COVID-19 pandemic as well as past pandemics and epidemics, scientists have focused on immunological studies to develop better vaccines as well as understand immune responses. Many of the questions are centered on studying T-cell epitopes, and peptide sequences that can be presented to immune cells to elicit responses against pathogens. Although current software can produce hundreds of predictions, they are generally not user-friendly nor graphical. In order to remove the existing programming barrier, we developed a Web tool to allow scientists to analyze and filter T-cell epitopes in an easy, intuitive, interactive, and versatile way. We have included two biological cases identifying new biological insights and showing the importance of having this type of toolset, especially for nonprogrammer researchers in the immunology field.

## Introduction

T-cell epitopes are peptides derived from processed antigens that are recognized by T-cells to elicit adaptive immune responses. Generally, CD4+ T-cells recognize epitopes between 13-17 amino acidic residues presented on the surface of major histocompatibility complex (MHC) class II molecules, while CD8+ T-cells recognize peptides of around 9 amino acid residues presented on the surface of MHC class I molecules (1). This allows T-cells to detect pathogens and abnormal self-antigens from cancer cells. Epitopes can be used to detect the magnitude of epitope-specific T-cell responses in an input sample based on cytokine secretion assays such as ELISPOTs or ELISAs (2, 3). Furthermore, the detection of epitope-specific T-cells has been used in diagnostic applications and to deimmunize proteins used as biological drugs (4, 5). Additional interest in T-cell epitopes is related to the cancer immunotherapy field, where the number of potential T-cell neoepitopes in a tumor has been proposed as a marker of success for checkpoint blockade treatments, and where tumor-specific epitopes are being used to induce tumor-specific T-cell responses (6).

The Immune Epitope Database and Analysis (IEDB) resource is a comprehensive database of experimental epitopes derived from individual low-throughput and high-throughput studies (7). There are three common categories of assays to identify T-cell epitopes considered by the IEDB. The first is assays measuring MHC binding in vitro to determine which peptides could be presented to T-cells (8). The second assay is MHC ligand elution where ligands are identified by mass spectrometry (9). The last assay consists in measuring T-cell response after the recognition of an epitope (10).

Given the complexity of experimental assays to measure epitope binding, several tools that predict epitope binding to different MHC alleles based on epitope amino acid sequences have been developed. Predictors such as SYFPEITHI (11) and BIMAS (12) use matrix-based methods making them fast algorithms but not the most accurate. More recent epitope predictors are based on machine learning algorithms trained with experimental data. These software such as NetMHC (13), NetMHCpan and NetMHCIIpan (14), and MHCFlurry (15) have shown to over-perform first generation methods. T-cell predictors provide a raw score (predicted binding score), and percentile rank (relative binding affinity) for each peptide, being percentile ranks the most widely used metric to filter T-cell epitopes (16).

These predictors often return a large number of predicted T-cell epitopes which need to be further filtered before experimental testing. Epitope features such as the predicted binding strength, the promiscuity (i.e., the number of MHC alleles they could bind to), the conservation across homologs, and the location within the amino acid sequence of the antigens (**Figure 1A-C**) allow an adequate filtering and selection process. However, there are few software automatizing this process. EpitopeViewer (17) is a Java application for the visualization of immune epitopes in IEDB allowing the identification of epitopes within the 3D structure of antigens or immunological complexes; however, this application does not show information about promiscuity or conservation of epitopes, and it is not currently maintained. IEDB allows analyses such as population coverage, epitope conservancy, and other tools such as cluster analysis or mapping mimotopes to antigens (18). However, most of these methods are not graphical and require some degree of programming experience, making them less accessible to non-programming researchers. Due to these shortcomings, we developed a user-friendly Shiny app named “Epitope-Evaluator”. This tool enables the analysis of T-cell epitopes such as the identification of conserved epitopes, promiscuous epitopes, or epitope-enriched protein regions focusing on the graphical interface; and facilitating its use for non-programming researchers.

**Figure 1.**
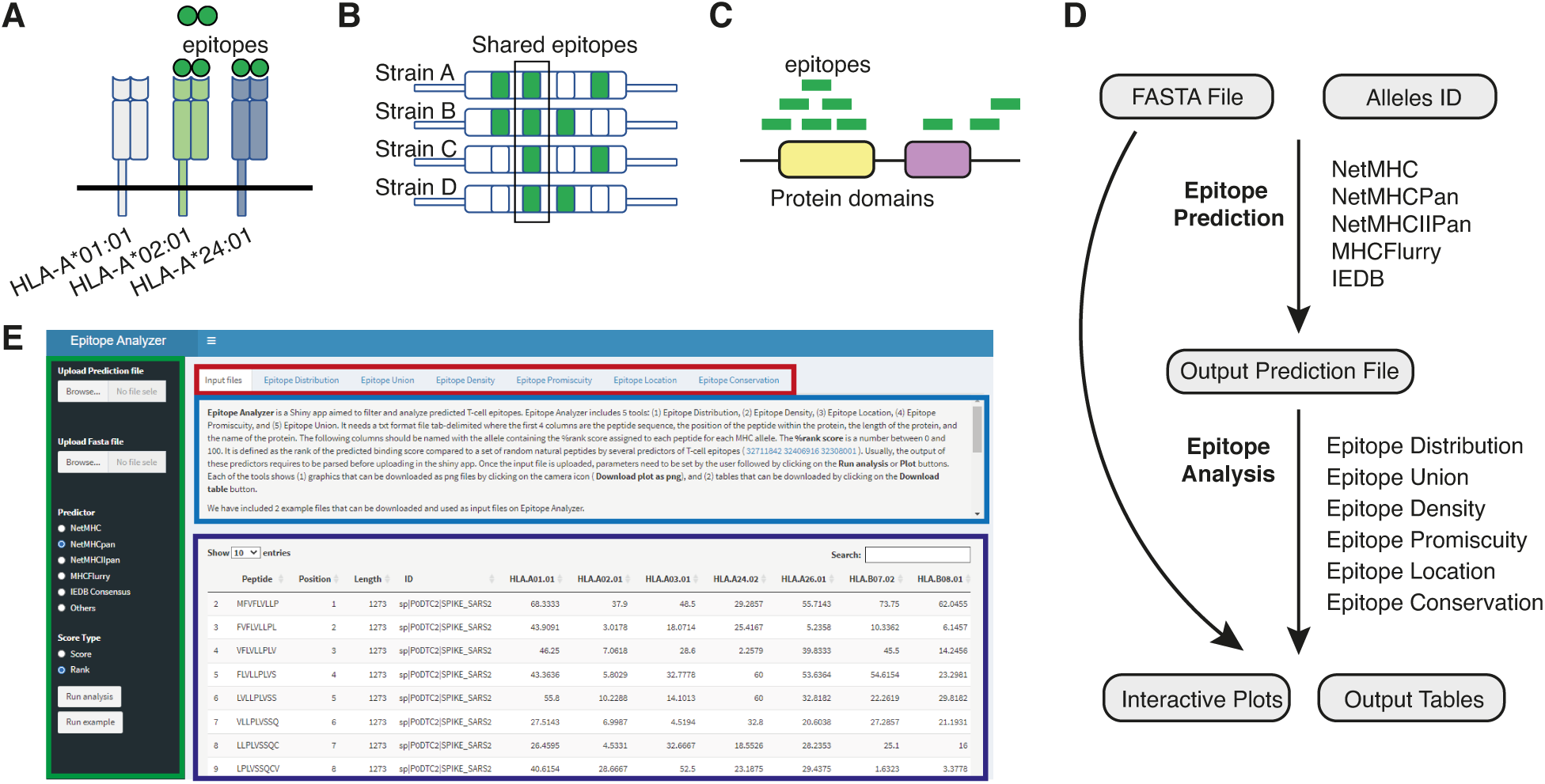
Schematic representation of characteristics of predicted epitopes and workflow of Epitope-Evaluator. **(A)** The promiscuity of epitopes represents their ability to be recognized by multiple Class I or Class II alleles. **(B)** The conservation of epitopes allows appropriate immune responses against several pathogen strains. Otherwise, restricted epitopes allow better discrimination across strains such as in immunological tests. **(C)** Determining the regions or domains enriched or depleted in epitopes helps to propose better peptide-based vaccines. Green circles and rectangles represent predicted epitopes. **(D)** Workflow to predict and analyze T-cell epitopes. All the current predictors of T-cell epitopes require a FASTA file containing protein sequences and the selection of MHC alleles of interest and retrieve an output prediction file containing a score for each epitope-allele pair (top). The Epitope-Evaluator requires the same FASTA file used as input in the prediction and the output file given by any predictor, and returns different plots and tables depending on the tool used (bottom). **(E)** Preview of the Epitope-Evaluator online web server. Rectangles indicate each section of the webserver. The parameters section (green), the title section (red), the help section (blue), and the output section (purple).

### Design and Implementation

Epitope-Evaluator is implemented in R v4.05 using the Shiny library. Prediction files are mainly processed with the dplyr package, and interactive plots are generated with ggplot2 and plotly packages. However, the app also requires the following additional libraries: readr, grid, gridExtra, reshape, shiny dashboard, tidyselect, rlist and tibble.

Epitope-Evaluator requires as input: 1) a multi-FASTA file containing the IDs and the sequences of the antigens, and 2) the prediction file obtained from a T-cell epitope predictor (**Figure 1D**). The user must indicate the predictor being used from a set of options or indicate “others” if not obtained from any of the listed predictors. In this case, the prediction file should have the following columns: the peptide sequence, its position within the protein, the protein ID, the protein length, and subsequent columns corresponding to each of the MHC alleles evaluated, where each value indicates a score for each epitope (**Figure 1E**). In addition to this, users must indicate whether the score in the table corresponds to the “percentile rank” or “binding affinity score”. The applicative will automatically identify whether the epitopes are class I or class II based on the name of the MHC alleles.

The title section indicates the different tools that are available in the web tool. Each of these tools is independent, thus, users can run all analyses in parallel. Each of the six tools and the ‘Input’ tab has four different sections: 1) The parameters section, 2) the title section, 3) the help section, and 4) the output section (**Figure 1E**). The parameter section, located on the left side of the web application, allows users to set different options and parameters for the corresponding tool. The help section describes the functionality of each tool, details each parameter, and explains the plots and tables returned in the output section. The output section shows the plots and tables from the selected analyses which are downloadable. All the tools contain interactive plots where users can zoom in/out, select regions, and obtain more information by hovering over the plots.

### Description of the Epitope-Evaluator Tools

Epitope-Evaluator facilitates the interpretation, visualization, and selection of predicted epitopes. This Shiny app can be used for analyzing MHC Class I and II epitopes and is compatible with any epitope predictor used. Epitope-Evaluator comprises six tools:

#### (1) Epitope Distribution

This tool helps to identify which MHC Class I or Class II alleles recognize the least number of epitopes and, therefore, which genotypes could be presenting a weaker T-cell response. The tool depicts a histogram representing the number of epitopes within a percentile rank or binding score range. This histogram can be shown per MHC allele or for the union or intersection of different MHC alleles, which can represent the number of epitopes that can be recognized by a heterozygote individual or the number of epitopes that are recognized by the alleles present in a population, respectively. Users can select between plotting a histogram (**Figure 2A**) or a cumulative histogram (**Figure 2B**). Hovering over any of the histogram bars provides the number of predicted epitopes with a percentile rank lower than a selected cutoff. In addition, the tool shows a heatmap indicating the number of epitopes predicted to bind to each allele with a percentage rank lower than a defined cutoff (**Figure 2C**).

**Figure 2.**
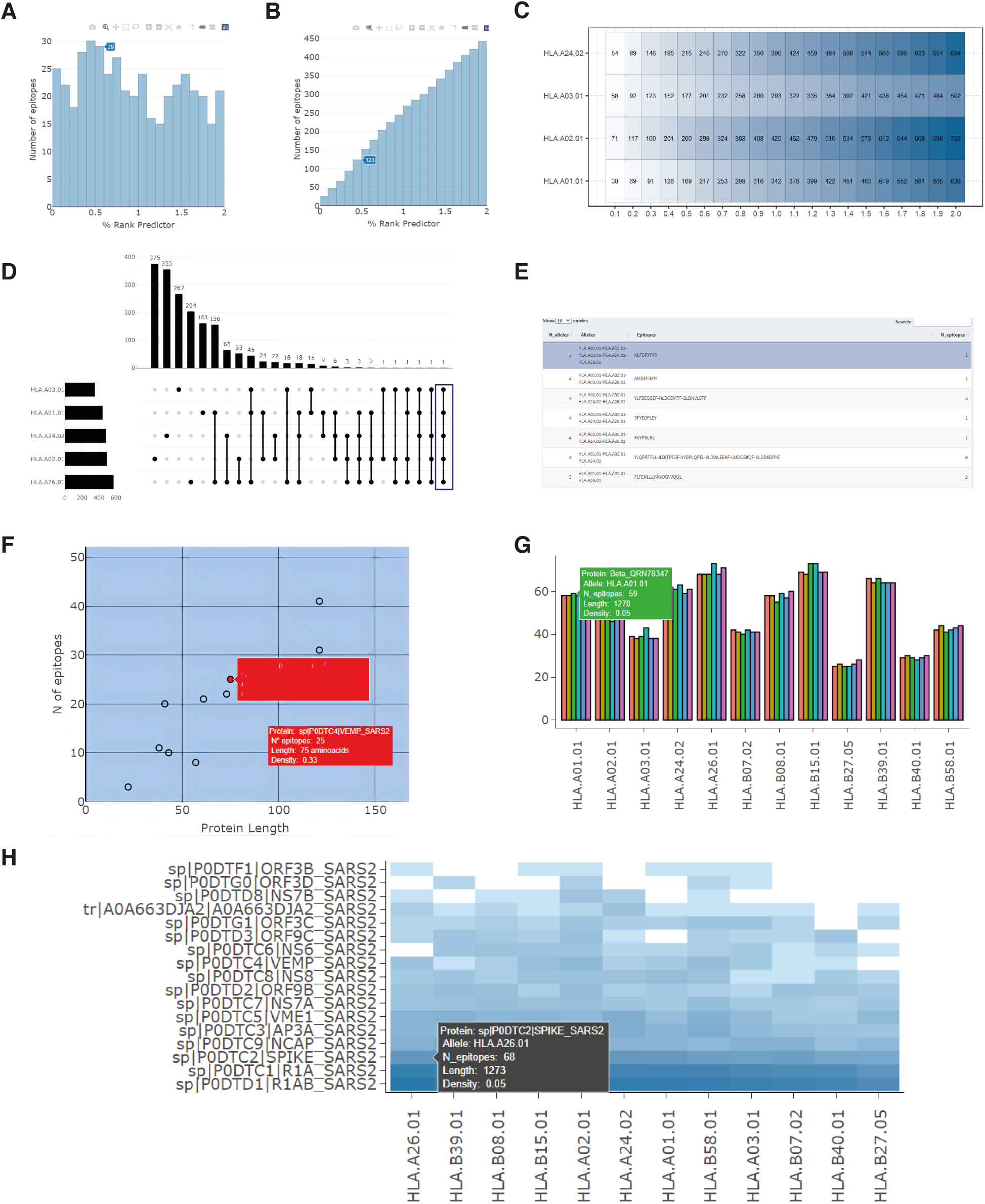
Plots produced with the (A-C) ‘Epitope Distribution’, (D-E) ‘Epitope Intersection’, and (F-H) ‘Epitope Density’ tools. **(A)** Histogram showing the number of epitopes within a particular range of percentage rank. **(B)** The cumulative histogram shows the number of epitopes with a percentage rank lower than a specified value. Hovering over any of the columns shows the corresponding number of epitopes. **(C)** The number of epitopes predicted to bind to each MHC allele considering different cutoffs. The cell color intensity represents the number of epitopes. **(D)** Up-Set plot, produced by the ‘Epitope Intersection’ tool, showing the number of epitopes shared by different combinations of five MHC Class I alleles. The five selected alleles are on the left side and the number of epitopes in each region is at the top. Individual points in the grid indicate epitopes binding to a specific MHC allele, while connected points indicate epitopes that can bind to multiple MHC alleles. **(E)** The table shows, for each region in the Up-Set Plot, the epitope sequences, the MHC alleles to which they are predicted to bind, and the number of epitopes within each region. **(F)** Scatter plot between protein length and the number of MHC Class I epitopes from SARS-CoV-2 proteins. **(G)** Bar plot showing the number of epitopes of five proteins variants predicted to bind to each MHC allele. The color of each bar represents a different protein variant. **(H)** Heatmap showing the number of epitopes within the SARS-CoV-2 proteins per MHC Class I allele. The cell color intensity represents the number of epitopes. Hovering over any point, bar, or cell (in the heatmap) shows more information such as the protein name, the number of epitopes, the MHC allele ID, the length of the protein, and the epitope density.

#### (2) Epitope Intersection

Different populations are represented by distinct combinations of MHC alleles. This tool enables us to identify the set of epitopes that could be used in epitope vaccines that are potentially recognized by all MHC alleles in the population, as well as to identify epitopes restricted to a particular set of MHC alleles. This tool shows the number of epitopes predicted to bind to different MHC allele combinations represented as a Venn Diagram or Up-Set plot if 6 or fewer MHC alleles are selected, or an Up-Set plot (26356912) if more than 6 MHC alleles are selected (**Figure 2D**). In addition to the downloadable plots, the tool provides a table containing the epitope sequences and the number of epitopes within each combination of MHC alleles (**Figure 2E**).

#### (3) Epitope Density

This tool can be used to determine the set of proteins containing a high number of predicted epitopes as a first step to finding highly immunogenic proteins. The tool displays a scatter plot of protein length versus the number of epitopes predicted to bind an MHC allele or combination of alleles (**Figure 2F**). The tool also shows the absolute number of epitopes within each protein predicted to bind to each MHC allele. This visualization can be displayed as a bar plot for a small number of proteins, (**Figure 2G)** or as a heatmap for several proteins (**Figure2H)**. In both cases, users can modify the plots by changing the fill range (i.e., by number or by the density of epitopes) and by arranging the set of proteins.

#### (4) Epitope Location

This tool can be used to identify the location of epitopes predicted to bind single and multiple MHC alleles, facilitating the visual identification of regions enriched with promiscuous epitopes. This tool graphically represents the position of each predicted epitope within each protein. Users must select a cutoff to identify epitopes and the MHC alleles of interest. Choosing the ‘Intersection’ option will show only epitopes predicted to bind to all of the selected MHC alleles. Selecting the ‘Union’ option displays all the epitopes which are colored ranging from yellow to red indicating the number of MHC alleles predicted to bind (**Figure 3A**).

**Figure 3.**
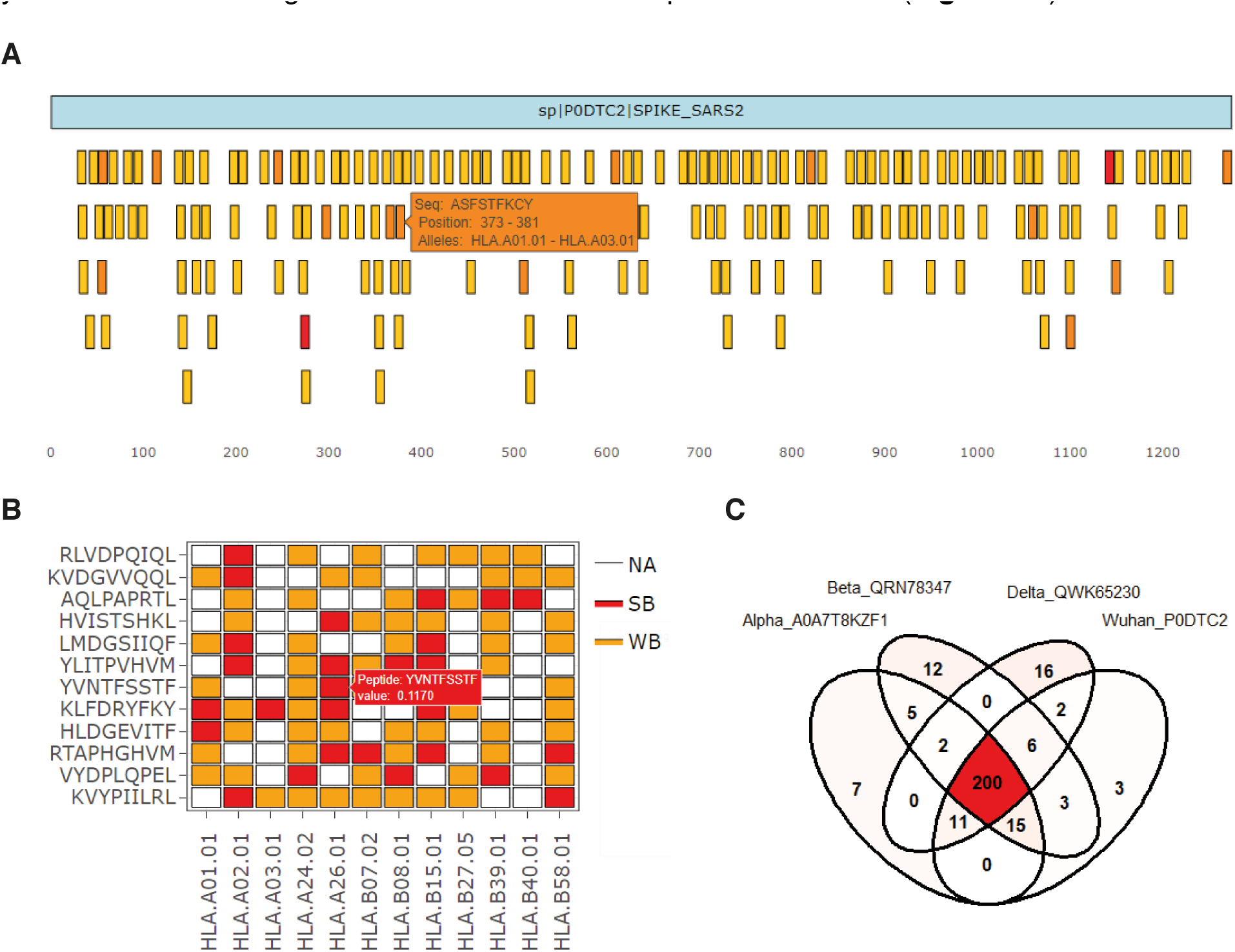
Plots produced with the *‘Epitope Location’, ‘Epitope Promiscuity’*, and *‘Epitope Conservation’* tools. **(A)** Linear representation of the SARS-CoV-2 Spike protein (light blue bar) and the MHC Class I epitopes (small bars). The color intensity of the represented epitopes varies from yellow to red indicating the number of MHC alleles predicted to bind to each epitope. Hovering over any epitope shows the sequence and the location of the epitope, and the MHC alleles to which the epitope is predicted to bind. **(B)** Heatmap showing the MHC Class I epitopes from the SARS-CoV-2 proteome that are predicted to bind at least seven MHC alleles. Red and yellow cells indicate strong (%rank < 0.5) and weak epitopes (%rank < 2), respectively. White cells indicate peptides that were not predicted as epitopes. **(C)** Venn diagram showing the shared epitopes found in the different Spike variants. The color intensity indicates the number of epitopes within each region.

#### (5) Epitope Promiscuity

This tool facilitates the identification of the promiscuous epitopes, which are epitopes predicted to bind to most MHC alleles, and their predicted percentage rank affinity. This tool shows in a heatmap the predicted epitopes that bind to more than a certain number of MHC alleles, set by the user. Users must also indicate the cutoff for both weak and strong binding epitopes. By default, these cutoffs are 0.5 and 2 for MHC Class I epitopes, and 2 and 10 for MHC Class II. The output of this tool is a heatmap where strong binder and weak binder epitopes are indicated as red and orange, respectively. Moreover, the tool returns a table with the sequence of the promiscuous epitopes, their start position within the protein, and the name of the corresponding protein (**Figure 3B**).

#### (6) Epitope Conservation

This tool allows users to identify epitopes that are present in multiple proteins, which can be useful to identify conserved epitopes across different pathogen strains. In addition, this tool allows for identifying epitopes gained or lost by diverse mutations. For this tool, users need to select the proteins, the MHC alleles of interest, and the cutoff percentile rank. The number of shared epitopes is represented as Venn Diagrams/Up-Set plot (≤ 6 proteins) or only Up-Set plot (> 6 proteins) (**Figure 3C**).

## Results

To illustrate the use of Epitope-Evaluator, we 1) analyzed the T-cell epitopes predicted from the SARS-CoV-2 proteome, 2) identified the most antigenic SARS-CoV-2 proteins and regions, and evaluated the density of epitopes across the Spike protein from different SARS-CoV-2 variants (**Supplementary Document 1**). First, we analyzed the epitopes of SARS-CoV-2 proteins across MHC Class I and II alleles. To identify which MHC alleles recognized the least number of epitopes from the SARS-CoV-2 proteome, and therefore may elicit a weaker adaptive immune response to infection, we used the “*Epitope Distribution*” tool. We found that, although most MHC Class I alleles recognize most epitopes, the HLA-B*27:05, HLA-B*40:01, and HLA-B*07:02 alleles showed the lowest number of predicted epitopes regardless of the percentile rank selected (**Figure 4A**). This suggests that patients with these alleles may have a weaker cytotoxic response against viral infection. Conversely, a similar number of predicted epitopes was found to bind each MHC Class II allele (**Figure 4B**), suggesting that variability in MHC Class II alleles would not be correlated with a worse or better viral response.

**Figure 4.**
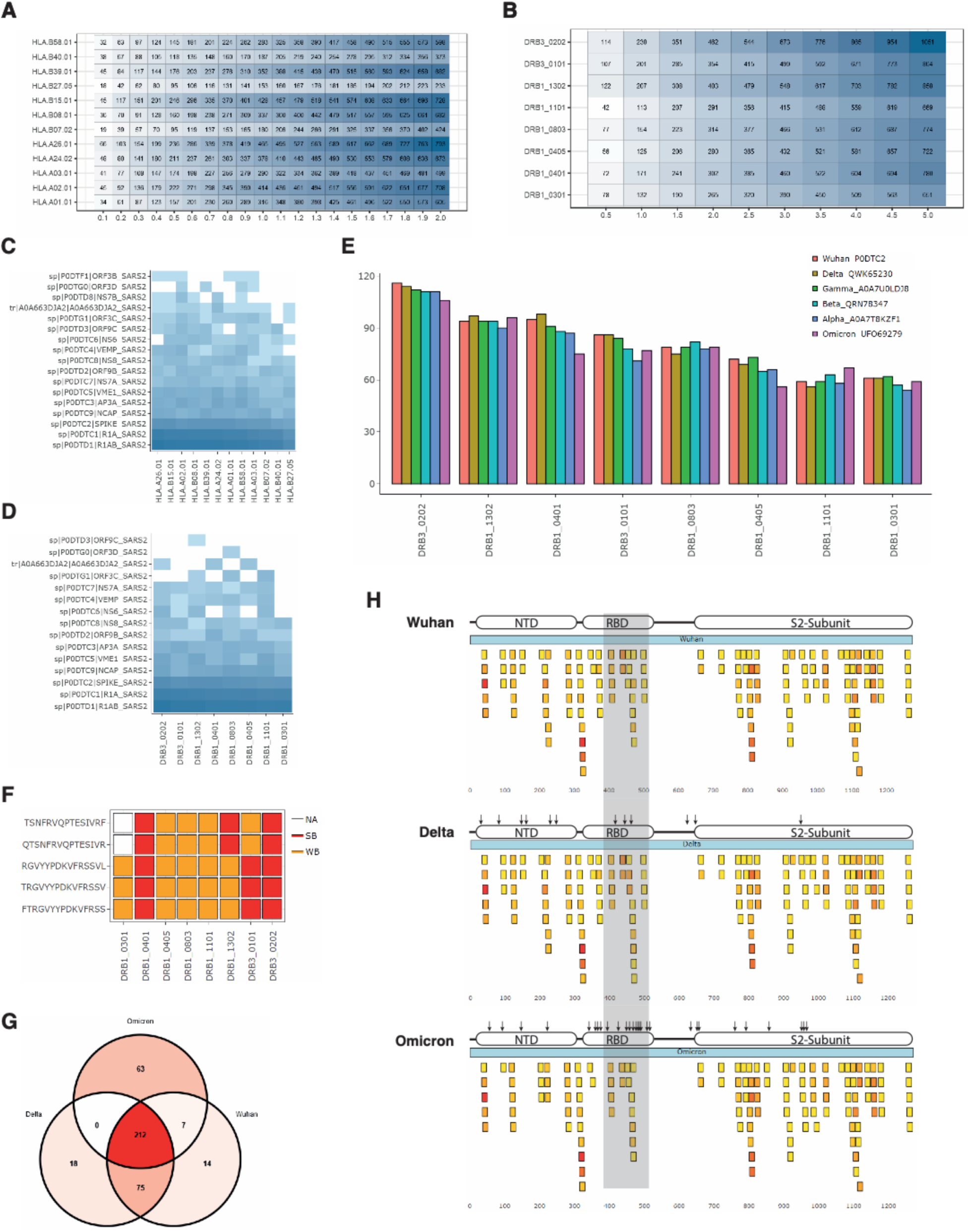
Analysis of MHC Class I and Class II epitopes from SARS-CoV-2 proteome and Spike variants. **(A-B)** Distributions of predicted epitopes across each **(A)** MHC class I and **(B)** class II alleles considering different percentage rank as cutoffs. **(C-D)** Heatmaps showing the number of epitopes in each protein predicted to bind to each **(C)** MHC class I or **(D)** MHC class II alleles. The color intensity represents a higher number of epitopes. **(E)** Bar plot showing the number of predicted epitopes within Wuhan Spike and its variants across eight MHC Class II alleles. **(F)** A heatmap representing the most promiscuous epitopes found in Spike protein. **(G)** Venn diagram showing the number of shared epitopes across Spike variants, and the number of gained and lost epitopes for Delta and Omicron variants. **(H)** Representation of Wuhan Spike, Delta Spike, and Omicron Spike, and their MHC Class II epitopes (%rank < 5). Spike proteins are shown as a light blue bar, while epitopes are represented as small bars colored from yellow to red representing the number of binding MHC alleles. A shadowed area is indicating the region where most. differences are located (amino acids 400 -500).

To identify which SARS-CoV-2 proteins are highly antigenic, we used “*Epitope Density*” and compared the number of epitopes in each protein per MHC allele. As expected, R1A and R1AB, which are the largest proteins, showed the highest number of both MHC Class I and II epitopes. From the remaining proteins, we show that Spike (S), Nucleocapsid (N) and Membrane (M) proteins contain the highest number of both MHC-Class I and II epitopes, suggesting the proteins N and M could also be considered complementary vaccine antigens, as previously suggested (19-23) (**Figure 4C** and **4D**).

To determine whether certain MHC Class II alleles recognize fewer epitopes from different Spike SARS-CoV-2 variants, we used the “Epitope Density” tool. We found that Wuhan Spike contains a similar number of epitopes compared to other Spike variants for each MHC Class II allele (**Figure 4E**). Further, we found that the five most promiscuous epitopes (epitopes binding to at least 6 MHC alleles), as determined using the *‘Epitope Promiscuity’* tool, were present in all six variants (**Figure 4F**). This suggests that these epitopes may be well suited for epitope vaccines as they are recognized by most MHC Class II alleles and are present in all current variants.

Next, we determined whether epitopes from Wuhan Spike are conserved in the Delta and Omicron variants, which could suggest cross-protection across variants when immunized against Wuhan Spike which is the antigen used in most current SARS-CoV-2 vaccines. Using the ‘Epitope Conservation’ tool we determined the number of epitopes shared, gained, and lost, by mutations in Delta and Omicron. We found 212 epitopes shared across the three variants, 21 and 18 epitopes were lost and gained in the Delta variant, and 89 and 63 epitopes were lost and gained in the Omicron variant, respectively (**Figure 4G**). This is consistent with the higher number of mutations in Omicron Spike compared to Delta Spike (24, 25). To determine whether the gain and loss of epitopes match with the mutation location, we used the ‘Epitope Location’ tool. We found that in the three variants, the region between amino acids 508 and 658 is depleted of epitopes. We also found that most differences between variants are located in the region between amino acids 400 and 500, which corresponds to the Receptor-binding motifs in the S1 region (**Figure 4H**). Altogether, our analyses suggest that although Spike variants can escape from certain antibodies elicited by vaccines against the Wuhan and Omicron Spike, the T-helper response may remain efficient, as demonstrated in other works (26, 27).

### Availability and Future Directions

Epitope-Evaluator represents a useful open-source program that allows the analysis and selection of candidate T-cell epitopes from different predictors. The biology application to evaluate SARS-CoV-2 epitopes demonstrates the relevance of having an integrative toolset to analyze epitopes in distinct contexts. The tool provides many customizable options and charts, reducing the computational skill barrier and making the analysis easier for non-programming users.

Epitope-Evaluator is available online at https://fuxmanlab.shinyapps.io/Epitope-Evaluator/. The application can also be run locally by downloading the code on its GitHub repository https://github.com/SotoLF/Epitope-Evaluator and launched locally in Windows, Mac OS, and Linux distributions. The example data are also included in the Web server so users can explore any of the tools. Moreover, we included a Rmarkdown file on GitHub showing each step of the biological application for reproducibility and transparency. We are releasing all the code under an open-source license and we promote collaborations and improvements from other researchers in the field. In addition to the Web application, the code may be easily dissected and implemented within individualized pipelines by other research groups.

Future directions include developing tools for analyzing B-cell epitopes and new ways to integrate MHC Class I T-cell epitopes, MHC Class II T-cell epitopes, and B-cell epitopes. Moreover, gene expression data for the antigens can be included to prioritize epitopes of highly expressed genes. Altogether, this tool will assist immunologists and experimental scientists to interpret and analyze T-cell epitopes.

## Acknowledgments

The authors would like to thank Manuel Ramirez and Luis Perez for their thoughtful comments during the development of this program.

## Author Contributions

**Conceptualization**

Luis Fernando Soto, David Requena.

**Data curation**

Luis Fernando Soto

**Formal analysis**

Luis Fernando Soto, Juan Fuxman Bass

**Funding acquisition**

Juan Fuxman Bass

**Methodology**

Luis Fernando Soto, Juan Fuxman Bass

**Project administration**

David Requena

**Resources**

Juan Fuxman Bass

**Software**

Luis Fernando Soto, Juan Fuxman Bass, David Requena

**Supervision**

Juan Fuxman Bass, David Requena **Validation:** Juan Fuxman Bass, David Requena **Visualization**

Luis Fernando Soto

**Writing/Editing**

Luis Fernando Soto, David Requena, Juan Fuxman Bass

## Supplementary Document 1

### Predictions of MHC Class I and Class II epitopes

We downloaded the proteome of SARS-CoV-2 from UNIPROT (ID: UP000464024) and used NetMHCPan 4.1 (1) and MHCflurry 2.0 (2) to predict 9-mer MHC Class I epitopes against supertype alleles: HLA-A*01:01, HLA-A*02:01, HLA-A*03:01, HLA-A*24:02, HLA-A*26:01, HLA-B*07:02, HLA-B*08:01, HLA-B*27:05, HLA-B*39:01, HLA-B*40:01, HLA-B*58:01, HLA-B*15:01.

The prediction in NetMHCPan was performed using the online server, while for MHCflurry we used the following command line: mhcflurry-predict-scan proteins.fasta --alleles HLA-A*01:01 HLA-A*02:01 HLA-A*03:01 HLA-A*24:02 HLA-A*26:01 HLA-B*07:02 HLA-B*08:01 HLA-B*27:05 HLA-B*39:01 HLA-B*40:01 HLA-B*58:01 HLA-B*15:01 --peptide-lengths 9 --no-throw --results-all--out Results_MHCFlurry.txt. To be conservative, we considered as MHC Class I predicted epitopes those peptides that showed a %rank less or equal than 2 in both predictors for the same MHC allele and assigned the highest %rank as its global %rank. A customized R script was used to filter both files and merge into one file to be used as input for Epitope-Evaluator. Similarly, we predicted 15-mer MHC Class II epitopes using NetMHCIIPanII 4.0 (1) considering a % rank cutoff of 5, and the following alleles DRB1-03:01, DRB1-04:01, DRB1-04:05, DRB1-08:03, DRB1-11:01, DRB1-13:02 and DRB3-01:01, and DRB3-02:02.

### Analysis of MHC Class I and II epitopes from the whole proteome of SARS-CoV-2

To run the Epitope-Evaluator with MHC-Class I predicted epitopes, we uploaded the input file and selected the option ‘Others’ as the predictor, since our file comes from a combination of two predictors, and the %rank as Score Type. Then we clicked on ‘Run Analysis’. We used the *Epitope Distribution* tool and used the following parameters. All alleles were selected, ‘Union’ was considered in Shared epitopes. The min %rank, max %rank, and the step were 0, 2, and 0.1, respectively. The Plot Type was ‘Cumulative Histogram’. We used the *Epitope Density* and selected 2 as Cutoff %Rank, Heatmap as Plot Type, Descendent as Sort Type, and Epitopes Number as Color by options. Then, we clicked on ‘Run analysis’ and ‘Generate plot’.

For MHC-Class II predicted epitopes, we uploaded the input file and the fasta file. We selected the ‘NetMHCIIpan’ as the predictor and ‘Rank’ as the Score Type. Then we clicked on ‘Run Analysis’. We used the *Epitope Distribution* tool and used the following parameters. All alleles were selected, ‘Union’ was considered in Shared epitopes. The min %rank, max %rank, and the step were 0, 5, and 0.5, respectively. The Plot Type was ‘Cumulative Histogram’. We used the *Epitope Density* and selected 5 as Cutoff %Rank, Heatmap as Plot Type, Descendent as Sort Type, and Epitopes Number as Color by options. Then, we clicked on ‘Run analysis’ and ‘Generate plot’.

### Prediction of MHC Class II epitopes from Spike variants

We downloaded the amino acid sequences of Spike variants from the NCBI using the following Genbank accession IDs (Alpha = QWE88920, Beta = QRN78347, Gamma = QVE55289, Delta = QWK65230, Omicron = UFO69279.1) and predicted the MHC Class II epitopes as in the previous analysis. We used NetMHCIIPanII 4.0 considering a % rank cutoff of 5, and the following alleles DRB1-03:01, DRB1-04:01, DRB1-04:05, DRB1-08:03, DRB1-11:01, DRB1-13:02 and DRB3-01:01, and DRB3-02:02.

### Analysis of MHC Class II epitopes from Spike variants

To run the Epitope-Evaluator, we uploaded the input file and the respective fasta file. We selected the ‘NetMHCIIpan’ as the predictor and ‘Rank’ as the Score Type. Then we clicked on ‘Run Analysis’. We used the *Epitope Density* and selected 5 as Cutoff %Rank, Bar plot as Plot Type, Descendent as Sort Type, and Epitopes Number as Color by options. Then, we clicked on ‘Run analysis’ and ‘Generate plot’. Then, we used the *Epitope Promiscuity* tool and selected 7 as the Minimum number of alleles, 1 and 5 as the Strong Binding Cutoff %rank, and the Weak Binding Cutoff %rank, respectively. Then, we clicked on ‘Run analysis’. To make the Venn Diagram, we used the *Epitope Conservation* tool. We selected ‘Wuhan, Delta, and Omicron’ as Proteins, 5 as Cutoff, all the alleles, and Venn Diagram as the Plot Type. Then, we clicked on ‘Run analysis’. Lastly, we used the *Epitope Location* tool. We selected the variant of interest (Wuhan, Delta, and Omicron), 5 as Cutoff %Rank, all the alleles, and union as ‘Shared epitopes. Then, we clicked on ‘Run analysis’

